# OmniPert: A Deep Learning Foundation Model for Predicting Responses to Genetic and Chemical Perturbations in Single Cancer Cells

**DOI:** 10.1101/2025.07.02.662744

**Authors:** Farzan Taj, Lincoln D. Stein

## Abstract

In cancer, intra- and inter-patient heterogeneity presents a significant challenge for therapeutic management, as patients with apparently similar profiles often exhibit divergent responses to the same therapies. This heterogeneity is primarily attributed to genetic and molecular variations among individuals and their tumors. Understanding the impact of these differences on treatment outcomes is widely believed to be a key step for developing effective precision medicine strategies. However, the complexity of most biological pathways makes it difficult to predict the effect of genetic variation on cells and tissues, let alone predict a patient’s response to therapy. As a result, high-throughput genetic and chemical perturbation screens have emerged as valuable tools for precision medicine-related tasks, such as disease modeling, target discovery, cellular programming, and pathway reconstruction. This approach is fundamentally limited, however, because the number of possible combinations of cell types, cell states, perturbation targets, and perturbation types is huge and cannot be exhaustively tested experimentally. This calls for computational approaches that can simulate such experiments *in silico*, guiding *in vitro* experiments towards perturbations that are more likely to produce the desired effect.

Here we describe OmniPert, a novel generative AI tool, which utilizes a deep learning, transformer-based architecture to model the effects of genetic and chemical perturbations on single-cell transcriptomes. Trained on millions of diverse cellular profiles, this approach allows for more granular analysis of cellular responses, thereby facilitating downstream applications in cell-specific gene-gene and gene-drug interaction networks, biomarker and drug target discovery, drug repurposing, and *in silico* perturbation reverse-engineering. In the context of oncology, OmniPert promises to facilitate the discovery of novel cell type- and state-specific targets, ultimately contributing to more effective and personalized cancer treatments.

## Introduction

A significant challenge for cancer care is that patients with similar demographics, tumor types, and medical histories often respond differently to the same therapeutic regimens. This inter-patient variability in therapeutic response is attributed to germline genetic differences among patients, acquired somatic variations in their tumors, and the epigenetic footprints of environmental and lifestyle factors (Menden et al., 2018). In addition, many cancers display high degrees of intra-tumoural heterogeneity in which the tumour harbors multiple genetically varied subclones. This intra-tumoural heterogeneity is a critical obstacle to effective treatment as it can lead to incomplete responses to therapy, enabling a subset of the tumor cell population to survive and drive disease recurrence (Burrell et al., 2013; Fisher et al., 2013). Over the past decade, substantial advancements in biological profiling have ushered in the ‘omics’ era, significantly advancing our understanding of many disease processes (Hertz & McLeod, 2013). Precision medicine aspires to harness this vast amount of molecular data, coupled with the patient’s medical history and genetics, to create customized therapeutic strategies that are highly effective with minimal adverse effects and a lower probability of relapse (Hertz & Rae, 2015). In pursuit of this vision, over the course of the last decade several consortia have undertaken large-scale systematic pharmacogenomic screening of panels of drugs across multiple molecularly profiled cancer models (Barretina et al., 2012; Garnett et al., 2012; Iorio et al., 2016; Yang et al., 2013). Despite advances in experimental automation and throughput, running such experiments on all possible combinations of compounds, cell lines, and genetic backgrounds is infeasible due to the rapid combinatorial increases in time and cost. Furthermore, only a fraction of the genome is directly targetable by existing drugs, meaning that many cancer targets remain inaccessible to direct therapeutic intervention and are missed by empirical drug screens (Sharma et al., 2024). These considerations call for a computational approach to predict drug efficacy in new cell types and genetic backgrounds.

To help address this gap, we recently developed and published an open-source, deep-learning algorithm called Multi-Modal Drug Response Predictor (MMDRP) (Taj and Stein 2024), which improves upon previous drug response prediction (DRP) approaches by overcoming several limitations in generalizability, data processing, and featurization. The use of new deep-learning methods and integration of multi-omic data significantly improved the prediction accuracy for novel drug and cell line combinations relative to models that learn from a single omic modality. However, a major limitation of MMDRP is that it was trained on bulk omic data derived from cell lines with potentially heterogeneous populations and included chemical perturbations only.

Bulk omic chemical screening experiments introduce two sources of ambiguity: First, bulk omic approaches obscure single-cell heterogeneity by capturing only averaged cellular responses, which can mask important cell-specific drug sensitivities or resistances. Second, the frequent absence of known drug target information reduces the model’s ability to capture broader molecular interactions, especially those involving untested compounds or novel drug-target relationships. Together, these factors significantly limit the predictive power and generalizability to diverse biological contexts, especially those involving the prediction of a heterogeneous tumor to therapy.

Recent advances in microfluidics, multiplexed barcoding, and sequencing have enabled precise profiling of transcriptomic responses to genetic and chemical perturbations at the single cell level. For almost a decade, this has allowed for large-scale studies that provide a higher-resolution view of heterogeneity in perturbation responses (Dixit et al. 2016; Adamson et al. 2016). The data produced by these single cell omics technologies enable the discovery of unique cellular states and responses often masked in traditional bulk population analyses, while encouraging the creation of more accurate predictive models based on individual cell type, cellular state and transcriptional networks.

In a typical single-cell perturbation study, scRNA-seq is performed on both pre-perturbed (control) and post-perturbed cells. Control cells are sequenced directly, while experimental cells are first exposed to a library of barcoded perturbations and then sequenced, enabling each unique perturbation to be linked to its transcriptomic response.. In particular, CRISPR-based perturbations allow us to target specific genes for overexpression or inhibition, either singly or in combination. This is in contrast to chemical perturbation screens where the definitive set of targets is not known ahead of time. Therefore, the integration of single-cell chemical and genetic perturbation modalities has the potential to better capture molecular interactions and address the aforementioned sources of ambiguity in perturbation modeling.

Here, we have extended work on MMDRP by developing an advanced transformer-based deep learning model called OmniPert that can jointly model the effect of both chemical and genetic perturbations of different kinds on single-cell transcriptomes. Besides predicting the post-perturbation transcriptional profile of various cells, this model has numerous other downstream applications in related tasks, such as biomarker discovery, target identification, drug repurposing, and gene regulatory network reconstruction. We envision future iterations of this tool becoming useful for selecting potential drug targets for large-scale drug discovery efforts, ultimately contributing to more effective cancer treatments, improving patient outcomes, and reducing healthcare costs.

## Results

We describe the design, training and validation of the model, followed by illustrations of its application.

### Model and Data Overview

Initially used for natural language processing (NLP), the transformer neural network architecture (Vaswani et al., 2017) has provided breakthroughs in many other data-intensive fields where the contextual interpretation of data is critical (Dosovitskiy et al., 2020; Zhang et al., 2023). The primary distinguishing feature of this architecture is that it allows the model to selectively and dynamically focus on pertinent input data segments through its attention mechanisms. This makes transformers a suitable architecture for modeling biological processes in which only a restricted set of genes are expressed in any given cell and cell state. **Figure 1** describes the OmniPert architecture, where we adapt the transformer architecture to single-cell perturbational data.

**Figure 1.**
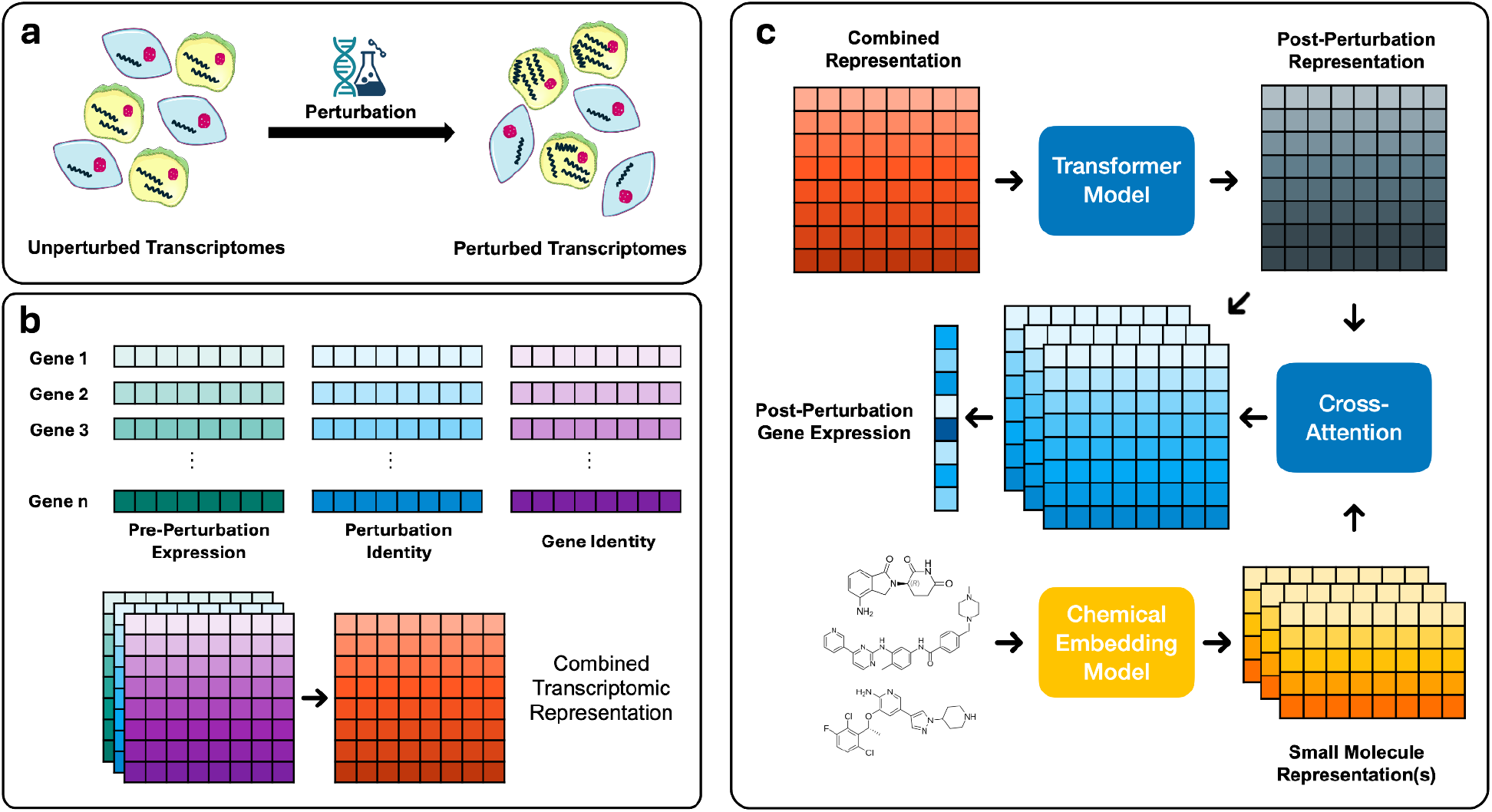
Simplified schematic of OmniPert model architecture. (a) Perturbations can change the transcriptional profile of some cells but not others. (b) Three types of information about each gene is used as the biological input: embedding of mRNA counts, or gene expression (cyan), embedding of gene perturbations and their types (blue), and embedding of gene identity (purple). These embedding representations for each gene are summed, resulting in a matrix of combined biological representations (red). (c) Chemical perturbations used in the sample is also represented using a fixed-size matrix (yellow).The transformer model processes the combined biological representations (red) into their embeddings (gray), followed by a cross-attention of drug and embedded genetic representations (yellow and gray, respectively). This results in a reconstructed mRNA count representation (blue), from which the predicted post-perturbation mRNA counts are extracted.

To obtain training and testing data, we used the scPerturb database (Peidli et al., 2024), a compendium of publicly available single-cell perturbation studies from a variety of tissue culture cell lines and patient samples. The studies comprising this database consist of the single cell RNA-seq (scRNA-seq) profiles of unperturbed cells, paired with profiles obtained after perturbation with drug(s) and/or genetic perturbation(s) via systematic CRISPR gene deletion, activation, and inhibition, as well as overexpression of selected ORFs. To assemble the dataset used for training and validation, we filtered and harmonized 20 single-cell perturbation studies from scPerturb based on whether they contained the above mentioned perturbation types paired with control data for each cell line (see Methods, **Tables 1 & 2**). This resulted in an aggregate dataset of over 6.25 million cells, including around 4 million genetically perturbed cells, 1.1 million chemically perturbed cells, 617 thousand cells with simultaneous genetic and chemical perturbations, and roughly 490 thousand control cells. After filtering, we used 5.2 million perturbed cells for training and 555 thousand perturbed cells for validation.

**Table 1.** Overview of datasets used in the making of the single-cell perturbation superset.

| # | Study Name | Cell Type / Line | Tissue | Number of Genes | Number of Cells | Number of Control Cells | Perturbation Type |
| --- | --- | --- | --- | --- | --- | --- | --- |
| 1 | Adamson - Weissman (2016) (Adamson et al., 2016) | K562 | Lymphoblast | 18,517 | 72,174 | 3,365 | CRISPRi |
| 2 | Aissa - Benevolenskaya (2021) (Aissa et al., 2021) | PC9 Xenograft | Lung Adenocarcinoma | 16,690 | 27,955 | 13,754 | Small Molecule Drugs |
| 3 | Chang - Ye (2022) (Chang et al., 2022) | PC9 | Lung Adenocarcinoma | 33,629 | 42,277 | 21,043 | Small Molecule Drugs |
| 4 | Datlinger - Bock (2021) (Datlinger et al., 2021) | Jurkat T-Cells | Immune | 15,265 | 13,383 | 1,107 | CRISPR-Cas9 |
| 5 | Dixit - Regev (2016) (Dixit et al., 2016) | K562 | Lymphoblast | 18,205 | 34,426 | 1,445 | CRISPR-Cas9 |
| 6 | Frangieh - Izar (2021) (Frangieh et al., 2021) | Melanocyte | Skin | 21,059 | 53,638 | 9,017 | CRISPR-Cas9 |
| 7 | Joung - Zhang (2024) (Joung et al., 2024) | hESC | Human Embryonic Stem Cell | 35,817 | 1,312,351 | 127,813 | ORF Overexpression |
| 8 | Lotfollahi - Theis (2023) (Lotfollahi et al., 2023) | A549 | Lung Epithelial Cells | 30,309 | 63,430 | 1,451 | Small Molecule Drugs |
| 9 | McFaline - Trapnell (2024) (McFaline-Figueroa et al., 2024) | A172, T98G, U87MG | Glioblastoma | 36,286 | 980,720 | 40,875 | Small Molecule Drugs, CRISPRi |
| 10 | McFarland - Tsherniak (2020) (McFarland et al., 2020) | 206 Cell Lines | - | 26,118 | 130,534 | 25,851 | Small Molecule Drugs, CRISPR-Cas9 |
| 11 | Norman - Weissman (2019) (Norman et al., 2019) | K562 | Lymphoblast | 20,128 | 111,445 | 11,855 | CRISPRa |
| 12 | Papalexi - Satija (2021) (Papalexi et al., 2021) | THP-1 | Monocyte | 17,593 (Normal) , 15,431 (Arrayed ) | 20,729 (Normal), 8,984 (Arrayed) | 2,386 (Normal), 2,009 (Arrayed) | CRISPR-Cas9 |
| 13a | Replogle - Weissman (K562 Essential) (2022) (Replogle et al., 2022) | K562 | Lymphoblast | 8,518 | 285,418 | 10,691 | CRISPRi |
| 13b | Replogle - Weissman (K562 GWPS) (2022) | K562 | Lymphoblast | 8,206 | 1,550,253 | 75,328 | CRISPRi |
| 13c | Replogle - Weissman (RPE1) (2022) | RPE1 | Retinal Pigment Epithelial | 8,695 | 214,190 | 11,485 | CRISPRi |
| 14 | Shifrut - Marson (2018) (Shifrut et al., 2018) | T-Cell | Immune | 16,231 | 22,591 | 13,461 | CRISPR-Cas9 |
| 15 | Srivatsan - Trapnell (2020) (Srivatsan et al., 2020) | MCF7, A549, K562 | Breast, lung, lymphoblast | 37,392 | 758,365 | 17,472 | Small Molecule Drugs |
| 16 | Tian - Kampmann (2019) (Tian et al., 2019) | Neuron, iPSC neuron | Neuron | 22,922 | 198,925 | 8,399 | CRISPRi |
| 17 | Tian - Kampmann (2021) (Tian et al., 2021) | iPSC neuron | Neuron | 22,123 | 53,493 | 871 | CRISPRi (32,300), CRISPRa (21,193) |
| 18 | Wessels - Satija (2023) (Wessels et al., 2023) | THP-1 | Monocyte | 15,104 | 30,707 | 424 | CRISPR-Cas13 |
| 19 | Xu - Cao (2024) (Xu et al., 2024) | HEK293 | Embryonic Kidney | 33,296 | 98,315 | 2,758 | CRISPRi |
| 20 | Zhao - Sims (2021) (Zhao et al., 2021) | Glioma brain samples | Glioma | 34,974 | 165,420 | 88,081 | Small Molecule Drugs |

**Table 2.** Overview of perturbation types in the single-cell perturbation superset.

| Perturbation Type | Number of Cells | Number of Unique Genetic Targets / Drugs |
| --- | --- | --- |
| CRISPR-Cas9 | 142,345 | 304 |
| CRISPRa | 120,349 | 201 |
| CRISPRi | 3,281,140 | 8,005 |
| CRISPR-Cas13 | 30,283 | 28 |
| ORF Overexpression | 1,184,538 | 1717 |
| Small Molecule Drugs | 1,167,973 | 211 drugs |
| CRISPRi + Small Molecule Drugs | 617,656 | 516 genes, 5 drugs |

The task for OmniPert is to predict the post-perturbation transcriptional profile of an individual cell given its pre-perturbation profile and the nature of the perturbation(s) (**Fig. 1a**). The model has two separate input heads: one for biological inputs, including the transcriptional profile and genetic perturbations, and another head for chemical perturbations. We fuse separate latent representations of gene identity, expression, and perturbation into a combined transcriptomic embedding (**Fig. 1b**). Chemical perturbations are embedded using Uni-Mol+ (Lu et al. 2023), chosen due to its robust performance on the Open Graph Benchmark (OGB) (Hu et al. 2020). For samples with a chemical perturbation, the latent chemical representations are integrated with biological data via a cross-attention layer that models the interaction of each drug with every expressed gene in the transcriptomic input, facilitating joint learning across genetic and chemical perturbations (**Fig. 1c**). As there is no single ground truth for unperturbed control cells, we choose a randomly sampled cell line and dataset-specific control cell for each unperturbed/perturbed example in the training and validation sets.

### OmniPert accurately simulates multi-modal perturbations in-silico

As the aggregated dataset encompasses a variety of perturbation types and combinations, we decided to apply a simplified 90:10 training and validation split based on unique perturbations stratified across cell lines and datasets to allow for equal representation across different conditions (see Methods). We then measured multiple metrics across perturbation types, cell types, and datasets. Without pre-training via e.g. self-supervised masked gene expression modeling, the model accurately predicts the post-perturbation profiles in unseen samples across all six major perturbation types tested, as measured with the Pearson Correlation between predictions and ground truth (**Fig. 2a**). We observe that gene pairs with stronger predicted perturbation effects are better correlated than those with smaller perturbation effects. Furthermore, the model captures both the magnitude and the direction of change across all genes, as well as those that are differentially expressed between the pre- and post-perturbation states (**Fig. 2b**). To assess directional accuracy, we binned log_2_-fold changes into three classes; up (Δ > 0.5), down (Δ < -0.5) and no-change (|Δ| ≤ 0.5), and constructed independent ROC curves for up- and down-regulation on the validation data. Discrimination was then quantified by the area under each curve (AUC). **Figure 2c** demonstrates that our model distinguishes true expression shifts from background with high fidelity in both directions. To complement distance-based metrics and evaluate the preservation of local topology, we computed the Local Inverse Simpson’s Index (LISI) (Korsunsky et al. 2019) at the level of each cell’s k-nearest-neighbor neighborhood (**Fig. 2e**). LISI quantifies the diversity of predicted and true cluster labels within each neighborhood, enabling us to assess how well the model recapitulates the fine-grained intermixing of cells induced by perturbations. We observed that a majority of clusters overlap between the two groups, but there are populations of outliers that do not, reflecting prediction errors in those populations. We also measured the distribution of Spearman Correlation Coefficients (SCC), to assess how non-linear trends are captured across different datasets (Supplementary Fig. 1). Most datasets have SCC medians between 0.7 and 0.8.

**Figure 2.**
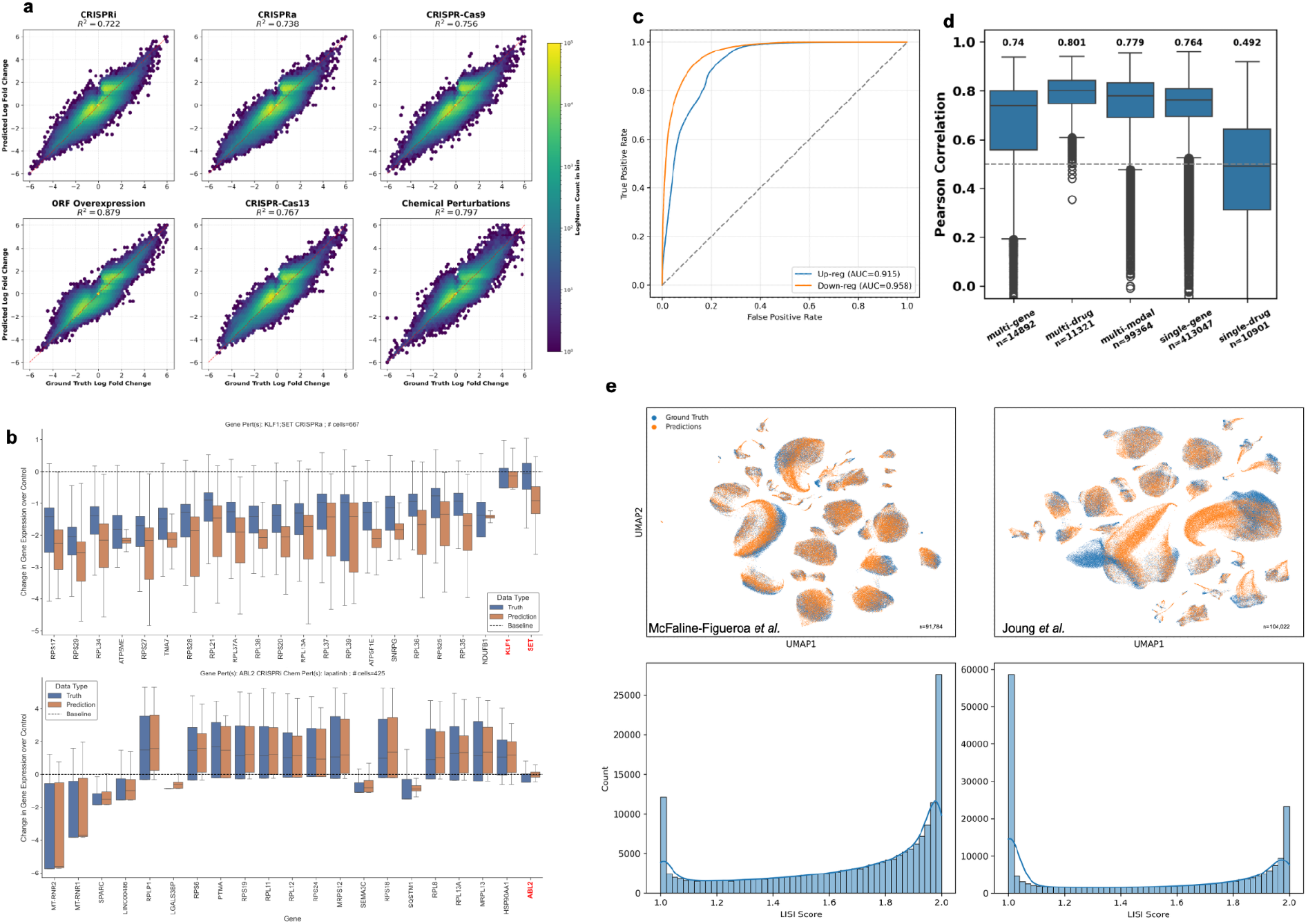
Overview of OmniPert’s validation performance. **(a)** Hexbin plots of post-perturbation Log Fold Changes (LogFC) relative to pre-perturbation; between predictions (y-axis) and ground truth (x-axis) in six genetic and chemical perturbation types. Coefficients of determination (R^2^) are reported above each panel. To make plotting feasible, we sampled 1e8 LogFC pairs from validation samples of each perturbation type. **(b)** Boxplots of two example conditions with combinatorial CRISPR-activation of two genes, KLF1 and SET (top) and CRISPR-interference of ABL2 combined with a chemical perturbation using Lapatinib (bottom). The boxes represent distributions of gene expressions relative to control (baseline at y=0). The chart is limited to the top 20 differentially expressed genes between pre- and post-perturbation, plus the perturbed gene(s) (highlighted in red). **(c)** ROC curves showing OmniPert’s performance in classifying true up-regulated (blue line) and down-regulated (orange line) gene expression changes, with respective AUC values indicating strong discriminative ability. Up- and down-regulation thresholds of 0.5 and -0.5 in log space, and subsample of 1 Million gene-samples for each curve were used. **(d)** Distribution of Pearson correlations among combinatorial and singular perturbations. Median Pearson correlations are denoted at the top of each box. Dashed line indicates a Pearson correlation of 0.5. **(e)** Top: UMAP representations of predicted and ground truth post-perturbation states from validation samples from two datasets, McFaline-Figueroa et al. (2024) (n=91,784) and Joung et al. (2024) (n=104,022). Bottom: Corresponding distributions of LISI scores for each dataset; higher is better.

To assess OmniPert’s performance on simultaneous perturbations, we subset validation samples into three combinatorial categories: multi-gene (samples with two gene perturbations), multi-drug (two chemical perturbations), and multi-modal (gene and drug perturbations) (**Fig. 2d**). In the multi-gene subset (n = 14,892), the median of per sample mean absolute error (MAE) was 0.435 and the median Pearson ρ = 0.740. The multi-drug subset (n = 11,321) achieved a median MAE of 0.370 and Pearson correlation ρ = 0.801, while the multi-modal subset (n = 99,364) yielded a median MAE of 0.339 and Pearson correlation of ρ = 0.780. Although performance on multi-gene combinatorial cases is modestly lower than on single perturbations (median ρ = 0.740 for multiple vs 0.764 for single perturbations), it’s slightly higher for multi-drug cases and in cases in which both genetic and chemical perturbations are applied.

Interestingly, predictions for single-drug perturbations (median Pearson ρ = 0.492) were substantially lower than for any combinatorial category. We hypothesize that this reflects the greater transcriptional heterogeneity inherent to single-compound assays, which in our validation set span over 200 distinct cell lines from the McFarland et al. (2020) study, each with its own baseline state and drug sensitivity. Each compound’s action likely engages cell-type-specific off-target and pharmacodynamic effects that OmniPert does not fully capture especially as genetic perturbation data for the same cell lines is not available. Together, these observations demonstrate that, although single-compound responses are inherently more variable, OmniPert robustly generalizes across increasingly complex perturbation scenarios.

### OmniPert helps identify gene and drug similarities

A recurrent issue in drug discovery and development is that a small molecule can have many targets, and it is often unclear which target is the biologically relevant one. We hypothesized that during OmniPert’s training on multi-modal data, it would learn biologically relevant similarities and differences among chemical and genetic perturbations in a way that can be used to clarify the target(s) of the chemical perturbation. To test this, we sought to interpret the trained model.

In OmniPert, interactions between genes are modeled via self-attention, while interactions between genes and small molecule drugs are modeled via cross-attention. The attention matrices from these mechanisms can be interpreted as a weighted adjacency matrix representing a network of either gene-gene or gene-drug interactions. We used the attention roll-out technique (Chefer et al., 2020) to interpret OmniPert’s attention matrices for a subset of the validation samples used as input.

We first selected chemically perturbed samples from the validation set whose predicted post-perturbation profiles correlated with the ground truth at Pearson r ≥ 0.8, considering both the full gene set and the subset of differentially expressed genes. Then, for each sample with a particular chemical perturbation, we generated an explanation matrix by performing a gradient-weighted integration of OmniPert’s attention matrices from multiple heads and layers into a single composite attention map (**Fig. 3a**). In the graph representation of this explanation matrix, we subsetted for nodes with the highest weighted degree centralities, essentially prioritizing genes the model attends to the most (**Fig. 3b**). Each drug perturbation has multiple samples and thus multiple graphs. We prioritized genes that appear the most often across these graphs, and made a gene-count weighted star graph for each chemical perturbation (**Fig. 3c**). The combination of graphs from all the selected chemical perturbations samples resulted in a bipartite graph where genes are only connected to drugs and vice versa. The bipartite projection of this graph reveals gene-gene and drug-drug relationships based on the number of shared drugs and genes, respectively (**Figs. 4a and 4b**).

**Figure 3.**
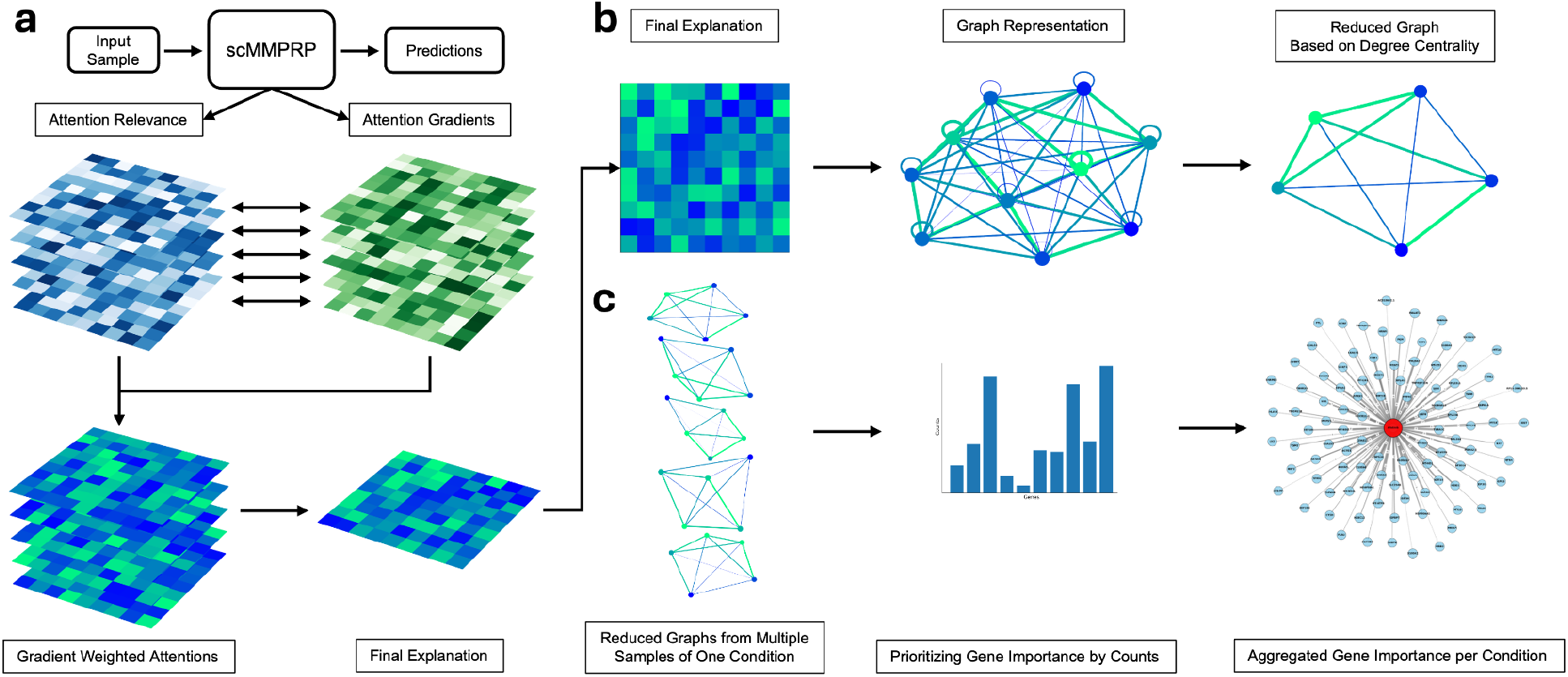
OmniPert Explanation Workflow. **(a)** Overview of attention roll-out and condition specific gene-importance graphs. The selected samples are passed to the model as inputs, and gradients adjust the attention weights, which are then multiplied into a final explanation. **(b)** The final explanation matrix is represented as an adjacency matrix, where self-connections are removed, and nodes are subset for the highest degree centrality, resulting in a reduced graph. **(c)** Reduced graphs from different samples of the same condition are pooled and genes are ranked based on the number of times they appear across reduced graphs of the same condition. The final result is represented as star graph for each condition, where the central node is the condition itself and the other nodes are genes associated with that condition.

**Figure 4.**
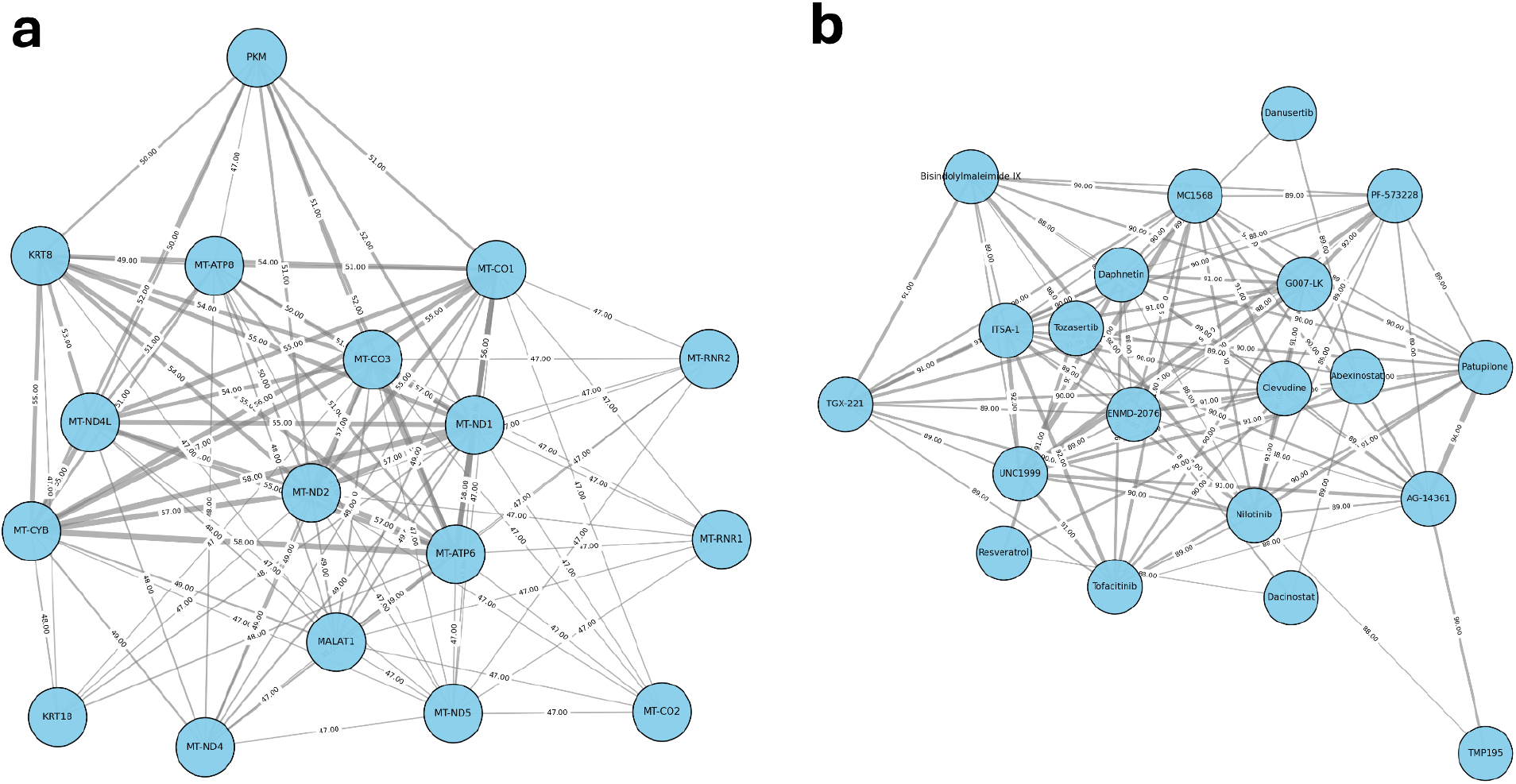
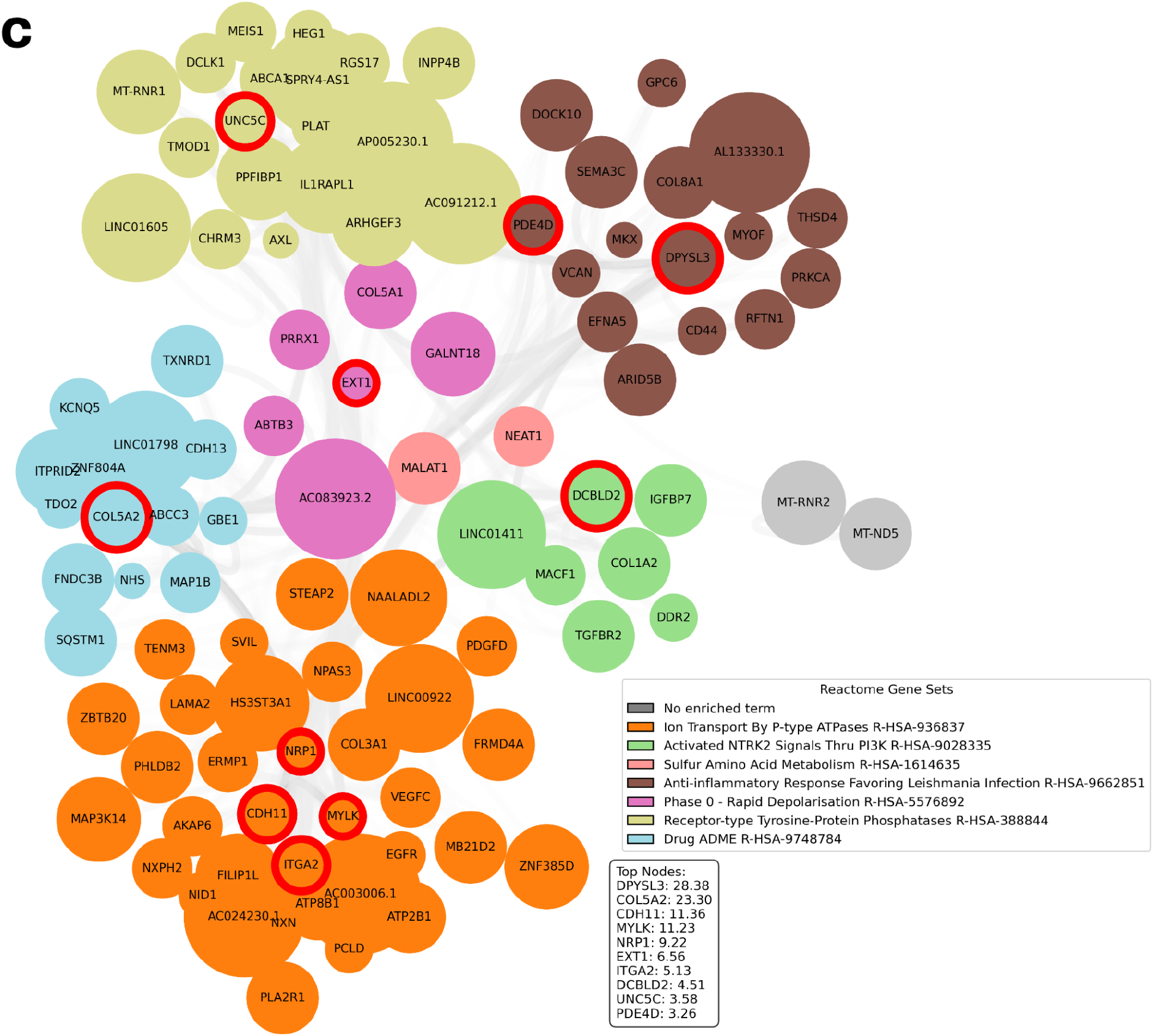
**(a)** Bipartite projections on genes and **(b)** drugs. We subset for top 100 edges in the star graphs of each chemical perturbation condition, and created two bipartite graphs based on shared drugs and shared genes (not shown). Bipartite projections of these two graphs hint at gene-gene and drug-drug relationships. **(c)** Graph of top gene-gene interactions in a sample from the A171 cell line from the McFaline-Figueroa et al. (2024) dataset, perturbed by MAPK7 CRISPRi genetic perturbation and Trametinib chemical perturbation. Genes are colored and grouped by Reactome gene set enrichment. Top nodes are listed and highlighted with a red circle. Certain pathways are highly interconnected, e.g., drug ADME (light blue) and ion transport (orange) gene-enriched subgraphs.

We assessed the top ten results from the drug bipartite projections, summarized in **Table 3**. These drug pairs are enriched with those that recapitulate known drug similarities and mechanisms of action, while others serve as hypotheses to further test in the lab. For example, in line with existing reports, the EZH2 inhibitor UNC1999 paired with the class IIa HDAC inhibitor MC1568 and with the FAK inhibitor PF-573228 both show confirmed synergy in preclinical cancer models. Additional pairs, such as Daphnetin + UNC1999 and Daphnetin + G007-LK, are mechanistically plausible. Daphnetin’s inhibition of JAK2/STAT3 and NF-κB signaling (Lv et al. 2024) could complement UNC1999’s relief of PRC2-mediated repression by jointly disabling inflammatory survival pathways and reactivating tumor suppressor and differentiation genes (Grinshtein et al. 2016); likewise, combining the tankyrase inhibitor G007-LK (which degrades β-catenin) (Voronkov et al. 2013; Mariotti et al. 2017) with Daphnetin’s blockade of cytokine-driven STAT3/NF-κB signaling may disrupt an autocrine/paracrine circuit that sustains Wnt-dependent cancer stem-like phenotypes. We believe that further interpretation analyses can help drug discovery efforts by identifying previously unknown drug-target interactions or biomarkers of response to therapy.

**Table 3.** Top drug-drug similarities based on shared high-attention genes.

| Rank | Drug Pair | Known Interaction/Synergy |
| --- | --- | --- |
| 1 | **UNC1999 - MC1568**<br>(z-score: 14.99) | **Yes:** Both are **epigenetic modulators** (EZH2 methylation vs HDAC deacetylation); reverse gene silencing in complementary ways. (Lue et al. 2019) |
| 2 | **AG-14361 - Patupilone**<br>(z-score: 15.69) | **Partial evidence:** PARP inhibitors + microtubule stabilizers show enhanced efficacy (e.g. olaparib+taxane in TNBC). Not specifically reported for AG-14361+Patupilone, but expected. Both induce DNA damage and p53 pathway activation. (Lu et al. 2018) |
| 3 | **G007-LK - UNC1999**<br>(z-score: 14.78) | **Not reported:** Conceptually synergistic by targeting cancer stemness from two angles (Wnt and PRC2). No known antagonism. Both force differentiation and loss of self-renewal. |
| 4 | **G007-LK - PF-573228**<br>(z-score: 14.83) | **Yes:** Preclinical synergy observed. Hits both intrinsic growth (Wnt/β-catenin) and extrinsic survival/motility (FAK/integrin). Wnt and FAK co-inhibition dramatically reduces proliferation. Pathway crosstalk suggests strong synergy (each blocks escape route of the other). (Wörthmüller and Rüegg 2020) |
| 5 | **Daphnetin - UNC1999**<br>(z-score: 14.20) | **Not reported:** Hypothesized to be beneficial. Daphnetin (multi-kinase inhibitor) could complement UNC1999's reactivation of silenced tumor suppressors; no known |
|  |  | conflict. Both impede survival pathways. |
| 6 | **ITSA-1 - UNC1999**<br>(z-score: 14.05) | **No (likely antagonistic):** ITSA-1 (lab tool) counters HDACi effects, so it would oppose the gene activation caused by UNC1999. No synergy data; not a viable combo. ITSA-1 could reduce UNC1999-driven transcriptional activation |
| 7 | **ITSA-1 - Tofacitinib**<br>(z-score: 15.29) | **Not reported:** No known interaction; ITSA-1 (HDAC activator) and Tofacitinib (JAK inhibitor) operate independently. Unlikely synergy; not studied. Both reduce transcription but via unrelated mechanisms. |
| 8 | **Bisindolylmaleimide IX - Nilotinib**<br>(z-score: 14.99) | **Partial evidence:** Not formally reported as combo, but BIM IX effective against BCR-ABL<sup>+</sup> leukemia including TKI-resistant (T315I) cases. Nilotinib blocks BCR-ABL oncoprotein signaling. This suggests that combination could cover both wild-type and resistant cells. Both trigger apoptosis in CML via different routes. (Zhang et al. 2016) |
| 9 | **UNC1999 - PF-573228**<br>(z-score: 14.23) | **Yes:** Demonstrated synergy in vitro. EZH2i + FAKi combination shows significantly greater tumor cell killing than either alone. Together disrupt EZH2-FAK pro-tumor feedback. (Bownes et al. 2021) |
| 10 | **G007-LK - Daphnetin**<br>(z-score: 14.76) | **Not reported:** Mechanistically plausible synergy by co-targeting Wnt/β-catenin and JAK/STAT3/NF-κB. Expected to cooperate in eliminating cancer stem-cell survival loops. Cuts off Wnt-IL-6/STAT3 positive feedback in tumors, forcing differentiation and removing external survival cues. (Campolo et al. 2024) |

To assess whether the model was capturing biologically meaningful relationships rather than merely memorizing correlations, we visualized gene-gene interaction networks and observed that highly attended gene pairs were significantly enriched for shared biological processes. As an illustrative example, we specifically examined genes that received top attention in samples subjected to MAPK7 CRISPRi combined with Trametinib treatment to see if they aligned with known pathway interactions (**Fig. 4c**).

Multiple top-scoring genes from the ERK5 (MAPK7) + MEK perturbation (by Trametinib) experiment map to known nodes in the ERK signaling network. For example, DPYSL3 (CRMP4) directly binds MEK to inhibit ERK1/2 activation (Du et al. 2022), and both UNC5C and EXT1 serve as brakes on MAPK output, whose loss amplifies ERK pathway signaling (Yuan et al. 2020). In contrast, NRP1 and ITGA2 promote robust Ras-ERK signaling through growth factor and matrix-integrin axes (Zhang et al. 2023). Meanwhile, the PDE4D enzyme is upregulated in MAPK-inhibited cells to restore ERK activity via cAMP suppression (Delyon et al. 2024). Lastly, we observe interconnectedness between drug ADME and ion transport enriched subgraphs. These confirmed relationships underscore that the model-identified genes indeed lie downstream of, or feed back into, the ERK5-ERK1/2 signaling pathways, consistent with their proposed roles in mediating or modulating the cellular response to combined ERK5 interference and MEK inhibition.

### OmniPert can recommend the perturbations needed to achieve a desired transcriptional state

Counterfactual explanations have recently emerged as a method in explainable AI to identify how the model input must change in order to observe a desired output. In the case of OmniPert, given the transcriptional profile of an input cell and its desired post-perturbation state, the counterfactuals would be a set of input perturbations that will produce the desired post-perturbation transcriptional profile. For example, this could be applied to the challenge of taking a drug that has undesirable toxicities, and finding less toxic drug candidates that phenocopy it. Counterfactual generation can also reveal interventions that promote the differentiation of cancer stem cells into non-replicative, differentiated cell types, similar to existing treatment strategies applied to some cancers such as neuroblastoma.

Following (Verma et al., 2020), we require that counterfactuals be feasible, i.e. correspond to valid, achievable interventions within the domain; minimal, i.e. involve the smallest possible changes to reach the counterfactual outcome; and diverse, i.e. offer multiple distinct alternatives that explore different directions in feature space. For genetic perturbations, we considered two alternative CF generation methods: 1) freeze all model weights and inputs except for the perturbation input and use backpropagation to solve for the perturbation necessary to produce the desired transcriptional state; and 2) use a brute force approach to apply all possible perturbations for each gene individually, followed by a ranking operation.

We chose the brute force approach as a suitable baseline. This has the benefit of producing multiple feasible and diverse CFs which can then be ranked by the closeness of the predicted transcriptional profile to the target profile. The disadvantage of this approach is its computational costs, which increase rapidly when there are multiple types of perturbations involved. For example, OmniPert inference will need to be run on the same cell 5,211 times in order to brute force its way through 1,000 expressed genes across five genetic perturbation types and 211 chemical perturbations.

We generated at least 1000 single-perturbation CFs for each dataset, and ranked them based on the Mean Squared Error (MSE) metric (**Fig 5a**). Recently published methods, such as scGPT (Cui et al. 2024), generate and rank counterfactual perturbations exclusively from the set of perturbations observed in their original (training) dataset, which we call the limited-world scenario. By restricting the candidate set, this setup drastically reduces the search space and artificially inflates performance; fewer tested perturbations increase the chance of CFs appearing near the top of the ranking. In contrast, the real-world scenario considers all expressed genes eligible for perturbations, making ranking more accurate but also more challenging, as CFs can now rank farther from the top. We report CF generation performance in both scenarios in **Figure 5b**. As expected, the performance in the limited-world setting is generally much better than the real-world scenario. As an example, the Replogle *et al*. (2022) genome-wide perturbation screen (GWPS) has tested for more than 2000 gene perturbations, which is much higher than other datasets, and thus has a closer CF performance to the real world scenario. In the real-world scenario, ground truth perturbation ranks differ widely between datasets.

**Figure 5.**
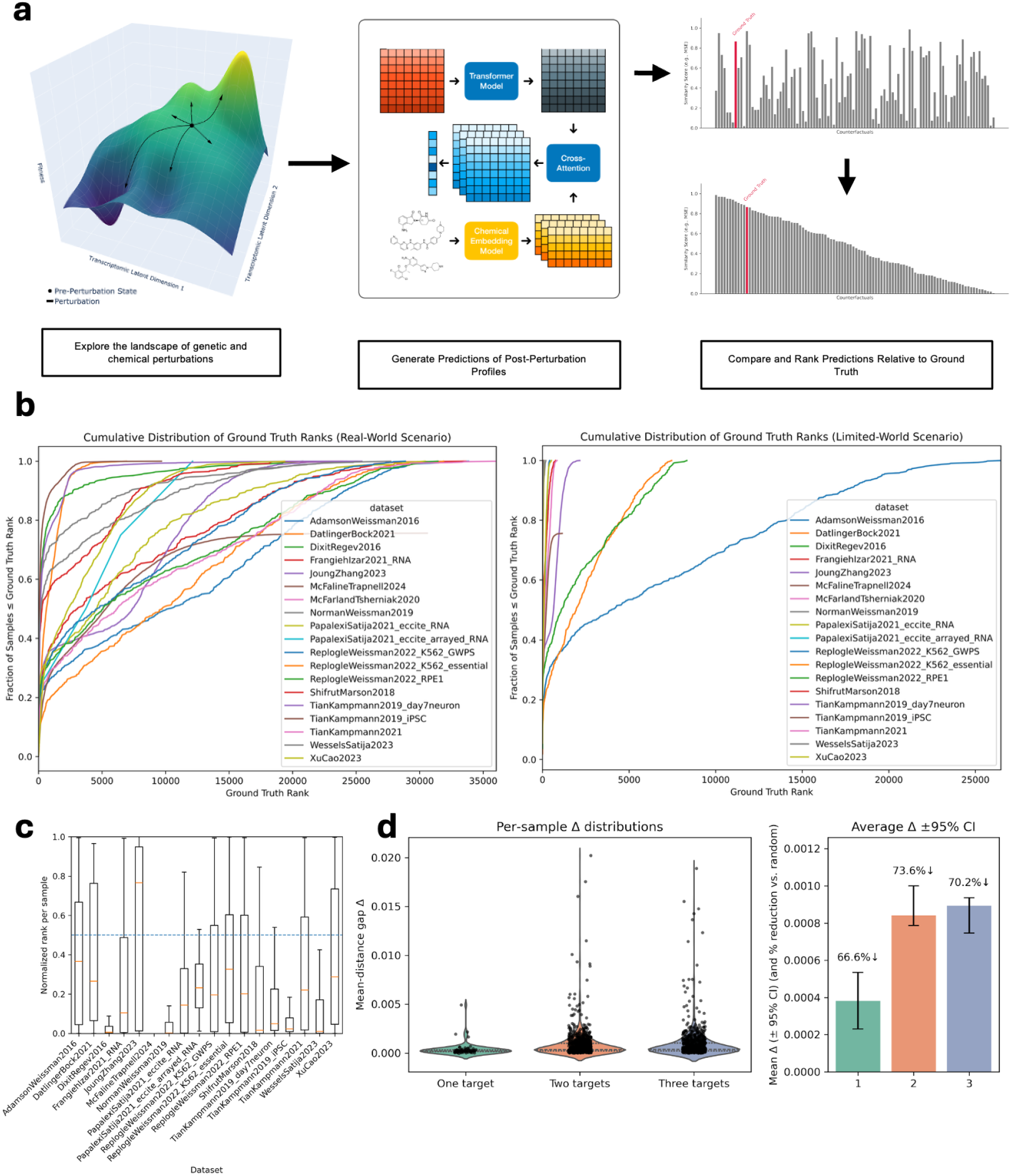
Counterfactual generation using OmniPert. **(a)** Overview of brute force CF generation and ranking. **(b)** Cumulative distributions of the ground truth ranks in the real-world and limited-world scenarios. The y-axis represents the fraction of samples whose ground-truth rank is less than or equal to the value on the x-axis. **(c)** Mean normalized counterfactual rank 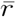 (± 95% CI) for each perturbation type, compared to the random expectation (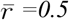 =0.5, dashed horizontal line). **(d)** OmniPert captures gene-drug perturbation concordance. (left) Distribution of per-sample cosine-distance gaps (Δ) between matching genetic and chemical perturbation targets versus random pairings, stratified by number of drug targets (one, two, three). Violin plots show density and medians (dashed), with individual sample values overlaid. (right) Mean Δ for each target category (bars) with 95 % CI (error bars) and the average percent reduction in distance relative to random perturbations (annotation). All Δ distributions are significantly greater than zero (Wilcoxon signed-rank test, p < 0.001), indicating that OmniPert embeds matching perturbations closer in the transcriptomic space.

Although we rank candidate counterfactual perturbations by the similarity between the model-predicted and the experimentally observed post-perturbation profiles, the ground-truth perturbation is not guaranteed to occupy rank one. Prediction error, transcriptional noise, metric choice, gene-subset masking, and genuine biological degeneracy can all cause an alternative perturbation to appear closer to the noisy empirical profile than the true one. Consequently, we report the CDF of ground-truth ranks, which captures how often the model succeeds despite these sources of discrepancy. We also calculated a normalized rank 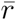 (see Methods) and compared its distribution relative to a random ranking baseline (**Fig 5c**). Across all samples tested for counterfactual generation, the average normalized rank was 0.29 (vs. random baseline of 0.50), indicating that empirically correct perturbations are on average in the top 29% of the candidate list.

To evaluate OmniPert’s ability to generate counterfactual perturbations that reflect its joint training on genetic and chemical data, we asked whether in silico CRISPR-based and small-molecule perturbations targeting the same gene produce concordant transcriptomic predictions. Specifically, we hypothesized that in-silico genetic perturbations (CRISPRi, CRISPR-Cas9, CRISPR-Cas13) and chemical perturbations directed at the same gene would yield closer predicted transcriptomic states than unrelated pairs. From the validation data, we selected a randomized subset of unperturbed cells across multiple cell lines, applied all single-gene and single-drug perturbations in silico, and computed pairwise cosine distances between the resulting post-perturbation expression profiles. Using a 10,000-replicate permutation test against random gene-drug pairings, we observed that matching pairs produced significantly smaller distances for one-, two- and three-target drugs, resulting in reductions of 66.6%, 73.6%, and 70.2% in distances relative to random gene pairings, respectively (Wilcoxon signed-rank test, p < 0.001) (**Fig 5d**, see Methods). These findings confirm that OmniPert embeds biologically meaningful concordance between genetic and chemical perturbations, supporting its utility for reverse-engineering drug mechanisms and target discovery.

## Discussion

Existing single-cell perturbation models, such as scGPT and CPA, have demonstrated the power of deep learning to predict transcriptional responses to either genetic or chemical perturbations in isolation, but they cannot readily handle both modalities or their combinations. OmniPert is, to our knowledge, the first transformer-based framework that simultaneously embeds genetic and small-molecule interventions at single-cell resolution and explicitly models their joint effects. By integrating self- and cross-attention over gene expression, gene identity, perturbation type, and drug embeddings, it can generalize to unseen perturbations, predict combinatorial responses, and generate counterfactual explanations. This advance moves beyond prior state-of-the-art by (i) unifying genetic and chemical perturbations in one architecture, (ii) supporting out-of-distribution inference for novel gene-drug combinations, and (iii) providing interpretable gene-gene and gene-drug interaction networks—thereby substantially broadening the toolkit for precision medicine applications.

Despite its strengths, OmniPert faces important limitations. Single-cell heterogeneity, both within and across datasets, complicates the pairing of control and perturbed cells. Future extensions could leverage clustering-based sampling strategies and incorporate domain-adaptive batch correction to mitigate these effects. Furthermore, while the analysis of genes that are perturbed with different technologies, such as CRISPRa versus CRISPRi, is useful for assessing the predictive power of perturbation models, different perturbation methods often come from different datasets, making it difficult to ascertain what proportion of the differences can be attributed to batch effects versus perturbation method.

Interestingly, ORF overexpression (with more than a million cells from the same dataset) achieves a markedly higher R^2^ (≈ 0.88) than the other five perturbation types (R^2^ ≈ 0.75). Because the ORF training data derive from a single experimental source, they have fewer batch-to-batch or lab-to-lab variations and often yield more uniform transcript-level changes, which likely makes them easier to predict. By contrast, the remaining perturbation modalities aggregate multiple datasets collected under different protocols and conditions, introducing additional noise and lowering overall predictive performance.

The reliance on multiple data sources warrants careful consideration of whether current and future perturbation datasets used for training such models should be integrated prior to training. One major concern with integration is that it risks diluting true signal by averaging across experiments with distinct technical biases, potentially masking subtle, perturbation-specific transcriptional changes. In addition, most of the currently available datasets do not contain samples in which a single cell is both genetically and chemically perturbed, or perturbed with multiple genetic or multiple chemical perturbations. Thus, the model’s performance in predicting such perturbation combinations cannot be as reliably assessed as singular perturbations. We hypothesized that the availability of genetic perturbation data would help offset the lack of large scale chemical perturbation data, by learning complementary underlying biological processes of perturbation response. However, expanding chemical libraries beyond the current 211 compounds used in OmniPert will still be essential to fully characterize combinatorial interactions and improve generalizability.

Some technical challenges also remain. First, the model’s assumption of a one-to-one correspondence between pre- and post-perturbation gene sets means that during training, we use the union of the two gene sets and set zeros for genes missing from either set. At inference time, we are forced to impute zeroes for genes silent in the pre-perturbation state. This limits the ability to predict de novo expression of genes that become activated after perturbation. If all possible gene inputs were to be included, training and inference would become computationally intractable. One option is to manually add genes of interest to the pre-perturbation state during inference. Overall, more research is needed in this area to allow for mismatching sets between the pre-and post-perturbation genes during inference. Second, we have demonstrated the potential for counterfactual generation and in-silico experimentation, but this process is highly sensitive to small prediction errors. Because counterfactuals often score closely on metrics like MSE, even minimal noise can change their ranking. This is compounded by intrinsic biological heterogeneity and experimental noise, and can substantially undermine the fidelity of in silico perturbations.

One promising route to improve both model predictions and counterfactual generation is to introduce a contrastive objective: for every cell we would push the model’s prediction for the true perturbation closer to the empirical profile while simultaneously pulling predictions for all other perturbations away. Such a loss would explicitly teach the network to maximise the margin between the ground truth and its nearest impostors, maximizing discriminability in latent space, while improving the counterfactual rank distribution in future iterations of the model. Lastly, by incorporating biological priors, such as pathway connectivity, cell-state annotations, and regulatory network structure, one could provide additional regularization to constrain counterfactual search and improve biological plausibility.

In the future, more sophisticated experimental paradigms, such as joint multi-modal perturbation assays, time-resolved single-cell profiling, and integrated spatial transcriptomics, will be critical to validate and refine *in silico* predictions. Expanding chemical libraries to include diverse compound classes, coupled with scalable CRISPR-based genetic screens, will enhance the model’s applicability across biological contexts. On the computational side, developing interoperable pipelines, benchmarking standards, and community-curated repositories will ensure reproducibility and facilitate method comparison. Although a substantial gap remains between predicting perturbations in individual cells and patient-level outcomes, these models can immediately support hypothesis generation for drug mode of action and in silico screening of compounds for desired cellular responses. Focused collaborations among experimentalists, modelers, and data scientists will be key to iteratively validate these predictions and drive near-term translational studies.

## Code and Data Availability

All the code and analysis scripts used for this project are publicly available on GitHub at https://github.com/LincolnSteinLab/OmniPert.

## Methods

### Model Inputs and Training Objective

The inputs of the model for the primary training objective are the pre-perturbation (control) transcriptional profiles, information on the perturbation target (in the case of genetic perturbations), and the type of perturbation. Each cell is treated as a single sample. For the latent representation of chemical perturbations, we use the Uni-Mol+ (Lu et al., 2023), a chemical property prediction model, as a ‘molecule embedder,’ selected for its performance on the Open Graph Benchmark (OGB) (Hu et al., 2020). The latent representation of small molecule drugs is combined with the latent biological representation through a cross-attention layer. The model predicts the post-perturbation transcriptional profile. We use the sum of two loss functions: Mean Squared Error (MSE) and a tanh (hyperbolic tangent) based direction loss. The main transformer module consists of three layers with eight heads each. The cross-attention module is a single layer with eight heads. The latent dimensionality across the model is 256.

We have programmed the model using the PyTorch (Paszke et al., 2019) and PyTorch Lighting (Falcon & The Pytorch Lightning Team, 2019) frameworks, which streamline model development, training, and validation and take advantage of GPU-based model training. The rest of the analysis is primarily performed in Python and associated packages.

### Training Data Preprocessing

We have created an aggregated dataset from 20 single-cell perturbation studies hosted by the scPerturb database (Peidli et al., 2024), which are summarized in **Table 1**. Each of these datasets is filtered to have a minimum of 100 genes per cell and a minimum of 10 cells per gene, followed by normalization and log transformation via Scanpy functions. After filtering out cells with unmatched controls, we limited transcriptome sizes to range between 100-5000 expressed genes. We set an upper bound of 5000 genes to balance between batch size, training speed and memory limitations. Because of this, out of the original 6.25 million cells, roughly 5.7 million cells remained. A perturbation-centric overview of the data is provided in **Table 2**.

**Supplementary Figure 1a** provides an overview of the distribution of transcriptome sizes across samples, which is not uniform, with many cells having too few expressed genes. Currently, samples with fewer than 100 and more than 5,000 expressed genes are filtered out.

Note that mismatches between gene- and transcript-naming conventions between the datasets results in a unique set of 77,500 unique gene identities. We converted all input genes from each dataset to a canonical HGNC symbol via synonym matching.

### Validation Split

We partitioned the dataset into training and validation subsets using a perturbation-centric 90:10 split, ensuring minimal data leakage while preserving model generalizability. Let *P* denote the set of all perturbations (genetic or chemical), *C* the set of cell lines, and *D* the set of experimental datasets. We randomly assign 10 % of the **unique** triplets (*p*,*c*,*d*) ∈ *P* × *C* × *D* to the validation set, with the remainder used for training. Under this scheme, a single perturbation *p*_*x*_ may appear in the training set in combination with other perturbations or in different contexts (e.g.,(*p*_*x*+y_,*c*,*d*) or (*p*_*x*_,*c′*,*d′*)), yet samples where *p*_*x*_ is the sole perturbation in a novel context are reserved for validation. This design challenges the model to predict out-of-distribution perturbations, such as new combinations, cell lines, or datasets, without substantial overlap between training and validation triplets, thereby limiting data leakage while testing true generalization.

### Identification of Differentially Expressed Genes in Validation Data

The validation data was divided into covariates based on both their cell line and dataset of origin. Each covariate, then has groups of cells, at the minimum the control cells with at least one other condition, e.g. a Cas9 deletion of a particular gene. All the cells from each covariate were then passed to scanpy function to rank gene groups, which uses the Wilcoxon rank-sum method to order genes based on importance in each group relative to the control group, effectively identifying differentially expressed genes.

## Model

### Biological Input Embeddings

Denote *n* by the number of genes in a cell and by *d* = 256 the embedding dimension of the transformer model. Let x = [*x*_1_,…,*x*_*n*_]^⊤^ ∈ ℝ^*n*^ be the original control expression values for *n* genes. We construct three biological embedding functions:

1. **Gene Expression Embeddings**. Each log normalized count *x*_*i*_ ∈ ℝ for gene *i* is via a two-layer multilayer perceptron comprising linear projections, SiLU activations, and layer normalization:

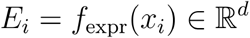

Masked or missing genes (indicated by a value of ™ 1) are embedded in the same space but do not contribute to training loss.
2. **Gene Perturbation Type Embedding**. We employ a learnable lookup table representing six modalities: CRISPRa activation, CRISPRi repression, CRISPR-Cas9 knockout, ORF overexpression, CRISPR-Cas13 knockdown, and an unperturbed state. In the future, newer modalities can be processed by expanding this lookup table. Each gene’s perturbation index *t*_*i*_ ∈ {0,…,*T*}is embedded via a learnable lookup:

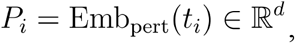

followed by layer normalization to ensure comparable scale.
3. **Gene Identity Embedding**. Each gene name *g*_*i*_ is first tokenized to an integer and then embedded:

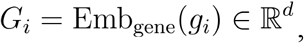

with optional initialization from a pretrained Gene2Vec matrix and a reserved padding index.

The combined **biological representation** for each gene is the element-wise sum:

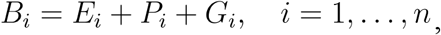

and stacking the *B*_*i*_ yields the matrix *B*. ∈ ℝ^*n × d*^

### Chemical Input Embeddings

Precomputed chemical embeddings for drugs are loaded from a precomputed lookup table where the UniMol+ model’s penultimate layer is treated as an embedding for each drug input. In effect, each unique SMILES string is mapped to a 200-dimensional vector which is stored in a lookup dictionary. These embeddings are further projected to match the transformer’s *d*-dimensional input space.

### Cross Attention and Output

For the samples where we have chemical perturbations, let *C*_*j*_ ∈ ℝ^*d*^ be the precomputed UniMol+ embedding for drug embedding index *j* = 1,…,*k*. Note that *k* is dependent on the number of atoms in each drug compound. We form the matrix *C* ∈ ℝ^*k × d*^ and compute cross-attention to integrate chemical context:

1. Project *Q = BW*_*Q*_,*K = CW*_*K*_, *V = CE*_*V*_,with attention matrices

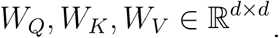
2. Compute attention weights

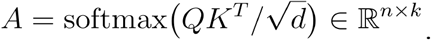
3. Obtain the chemical-conditioned update

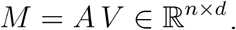

The final transformer input is the sum of biological and chemical streams:

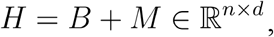

We recover a predicted expression vector of the same dimensionality via a simple linear read-out. Writing *w*_*out*_ ∈ ℝ^*d*^ for the learnable weight vector and *b* ∈ ℝ for the bias, each gene’s prediction is

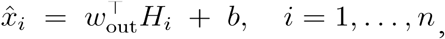

and in matrix form (note that *b*1_*n*_ is an *n*-vector full of *b*‘s):

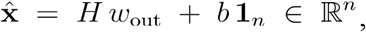

bringing us back into the original gene-expression space and enabling direct comparison to the input x via our loss function.

### Training Loss Objective

We trained OmniPert using a composite loss that balances accurate magnitude prediction with correct perturbation directionality. For each cell and gene, let x_pred_,,x_true_ and x_ctrl_ denote the predicted, ground-truth, and control expression values, respectively, and *m* be a binary mask indicating valid genes. The magnitude loss is a masked mean of the squared errors raised to the power 2 + γ, i.e.

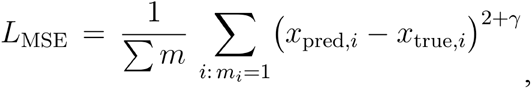

where γ = 2 penalizes large deviations more heavily and masking excludes padded or missing genes. The direction loss encourages agreement in expression change sign by comparing hyperbolic-tangent-transformed deltas,

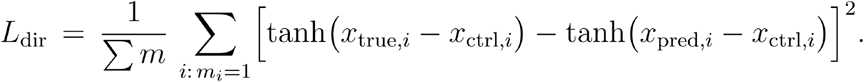

The total loss is a weighted sum,

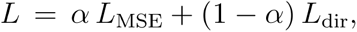

where we set *α* = 0.5 by default. This combination ensures the model not only minimizes prediction error but also captures the correct up- or down-regulation direction of each perturbation.

### Attention Roll-Out and Graph Generation

We interpret OmniPert’s learned self- and cross-attention as multi-modal interaction networks by applying the “attention roll-out” algorithm (Chefer et al. 2020) to a subset of high-confidence validation samples (Pearson ≥ 0.8). After registering backward hooks, for each selected sample, we performed a single forward pass to obtain each transformer layer’s raw attention tensor *A*^(*ℓ*)^, then executed a backward pass to retrieve the corresponding gradient tensor 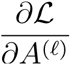, where *L* is the model loss.

**Head Aggregation**. Within each layer, we computed an element-wise Hadamard product of the attention and its gradient, clamped all negative values to zero, and averaged across attention heads to yield a non-negative, aggregated map *G*^(ℓ)^.

**Roll-out**. To integrate information across layers, we initialized a relevance matrix *R* as the identity matrix (*I*) and iteratively updated it according to

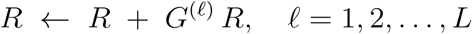

Each row of *R* represents the relevance of every token to the model’s prediction for that input. To construct gene-gene and gene-drug interaction graphs, we extracted the relevance profile for each perturbed token, retained the top *k* features by weighted degree for each sample, and aggregated these star graphs across samples of the same condition. These condition-specific graphs were then merged into a bipartite network connecting genes and drugs, and one-mode projections were computed to reveal gene-gene and drug-drug similarity networks. All graph operations were implemented in NetworkX and visualized using Matplotlib.

### Counterfactual Generation and Evaluation

We generated counterfactual perturbations for both genetic and chemical modalities by iterating through all single-gene and single-drug interventions for each unperturbed validation cell. First, we instantiated a custom dataset that, given a control cell, produces inputs for every candidate perturbation. We then batched inference via a data loader, passing each perturbation input through the trained OmniPert to obtain predicted post-perturbation transcriptomes. For each perturbation, we computed a set of similarity metrics (e.g., Spearman correlation) between the predicted profile and the ground-truth experimental profile; the resulting metrics were stored in a table and ranked. We recorded whether the true perturbation appeared in the top-K predictions as a primary performance measure, as well as the ranks of each counterfactual perturbation based on each metric. All results were saved per sample and organized by dataset and cell line for downstream analyses. Batch size, perturbation lists, and device placement were configured to maximize GPU utilization and minimize I/O overhead.

### Counterfactual Normalized Ranking

In order to quantify counterfactual (CF) accuracy in a single, comparable metric, we first compute for each sample *i* the normalized rank

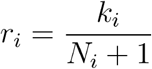

where *k*_*i*_ is the 1-based position of the true CF among *N*_*i*_ candidates. By construction 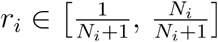 and a uniform random ranking yields E [*r*_*i*_] = 0.5. Samples are grouped by dataset *d*, and for each group we compute the mean normalized rank

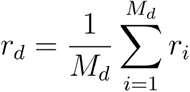

and its standard error

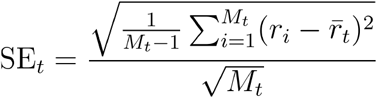

where *M*_*d*_ is the number of samples of dataset *d*. The resulting 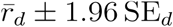 are plotted against the random baseline (*r* =0.5), and the full distribution of *r*_*i*_ is shown via boxplots to illustrate variation around the 0.5 expectation.

### Genetic and Chemical Perturbation Concordance in Counterfactuals

To evaluate whether OmniPert faithfully recapitulates known gene-drug relationships, we performed a concordance analysis on a randomized subset of unperturbed single-cell transcriptomes drawn from multiple cell lines. For each control cell, we generated in silico post-perturbation profiles by applying every single-gene inhibitory perturbation (CRISPRi, CRISPR-Cas9, CRISPR-Cas13) and every single-target drug perturbation using the trained OmniPert model. We then computed pairwise cosine distances between each gene-perturbed and drug-perturbed prediction, labeling pairs as “matching” when the drug’s known target gene(s) coincided with the genetic perturbation and as “random” otherwise. We obtained drug-target information from DGIDB (Cannon et al. 2024), and focused the analysis on low-promiscuity drugs with one, two or three annotated targets. For each sample, we summarized the concordance as the mean distance gap Δ (random minus matching) and expressed it as a percentage reduction relative to the mean random distance. Statistical significance of Δ > 0 was assessed by a one-sample Wilcoxon signed-rank test across samples, and results were visualized as per-sample Δ distributions (violin plots) and average Δ with 95 % confidence intervals and percent reduction annotations. This analysis demonstrates that matching perturbations are embedded significantly closer in transcriptomic space than non-matching controls.

### Drug-Drug Similarity Significance Testing

We began with a bipartite graph *G* whose two node sets are drugs and genes and whose edge weights *w*_*d*,*g*_ quantify the strength of each drug-gene association. To assess whether a given drug-drug edge (*d*_*i*_,*d*_*j*_) in the one-mode projection of *G* is stronger than expected by chance, we generated *R* = 1,000 random graphs {*G*^(*r*)^}each with 5000 edge-swaps “double-edge swaps” that each preserve every node’s degree and maintain bipartiteness. From each *G*^(*r*)^ we computed the weighted drug-projection and recorded the null weight 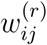 for the top 100 of the original edges. The empirical p-value for the observed weight 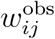 is

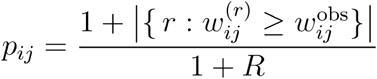

and the standardized score is

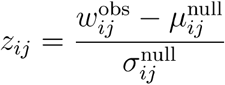

where 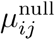 and 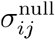 are the mean and standard deviation of 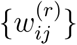. To control the false discovery rate across all tested pairs, raw p-values were adjusted by the Benjamini-Hochberg procedure (*α* = 0.05).

