## Supplementary Materials for "OmniPert: A Deep Learning Foundation Model for Predicting Responses to Genetic and Chemical Perturbations in Single Cancer Cells"

### scMMPRP Supplementary Materials

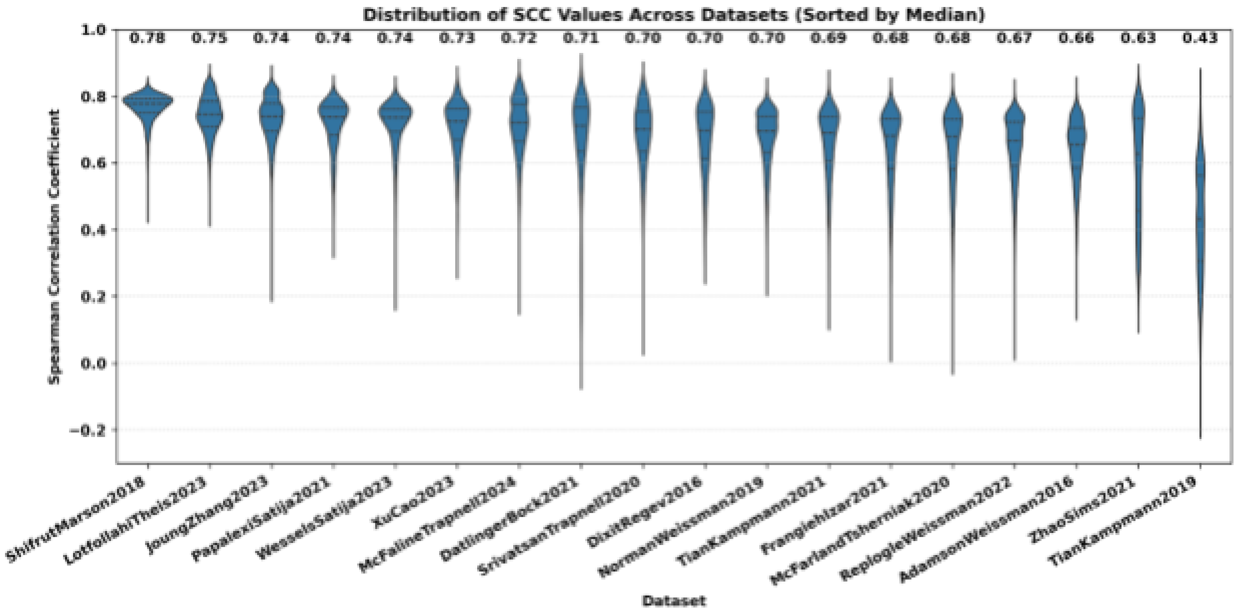


***Supplementary Figure 1.*** *Distribution of Spearman Correlation Coefficients (SCC) across datasets from the validation data.*

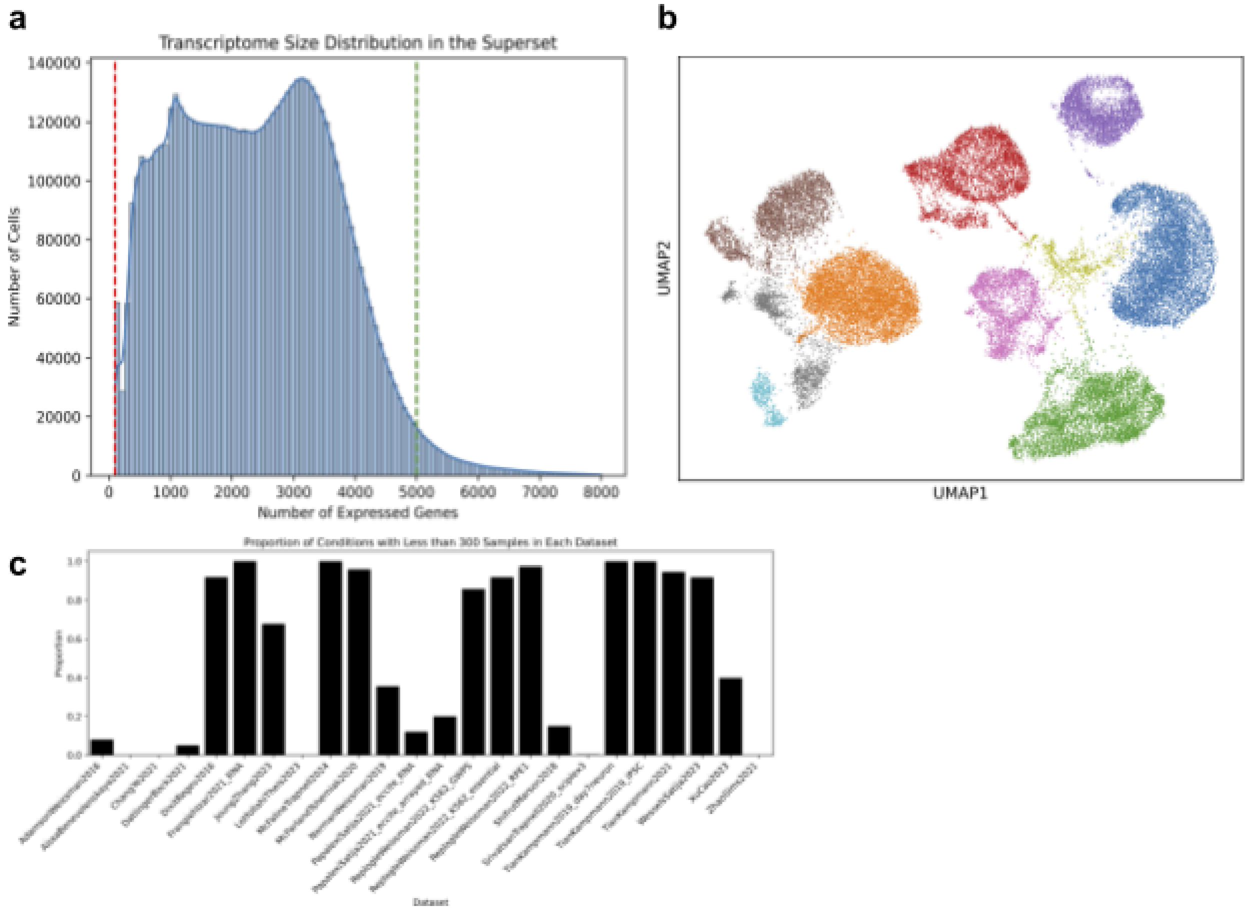


***Supplementary Figure 2.*** *Overview of certain superset data characteristics.* ***(a)*** *Distribution of transcriptome sizes across the unfiltered superset. As shown, the distribution is bimodal, with a large concentration around 100 and another around 3000 expressed genes. We have set lower and upper limits to 100 and 5000 expressed genes, as represented by red and green vertical lines, respectively.* ***(b)*** *UMAP of control samples from the K562 cell line aggregated from Adamson, Dixit, and Norman datasets, colored by Leiden clusters. The control samples display both inter- and intra-dataset heterogeneity, making pre-perturbation selection a non-trivial task.* ***(c)*** *Proportion of conditions with less than 300 samples in each dataset. To achieve sufficient statistical power, Peideli et al. suggested a minimum of 300 cells per condition. Having too few samples per unique condition may also play a role in poorer generalization.*

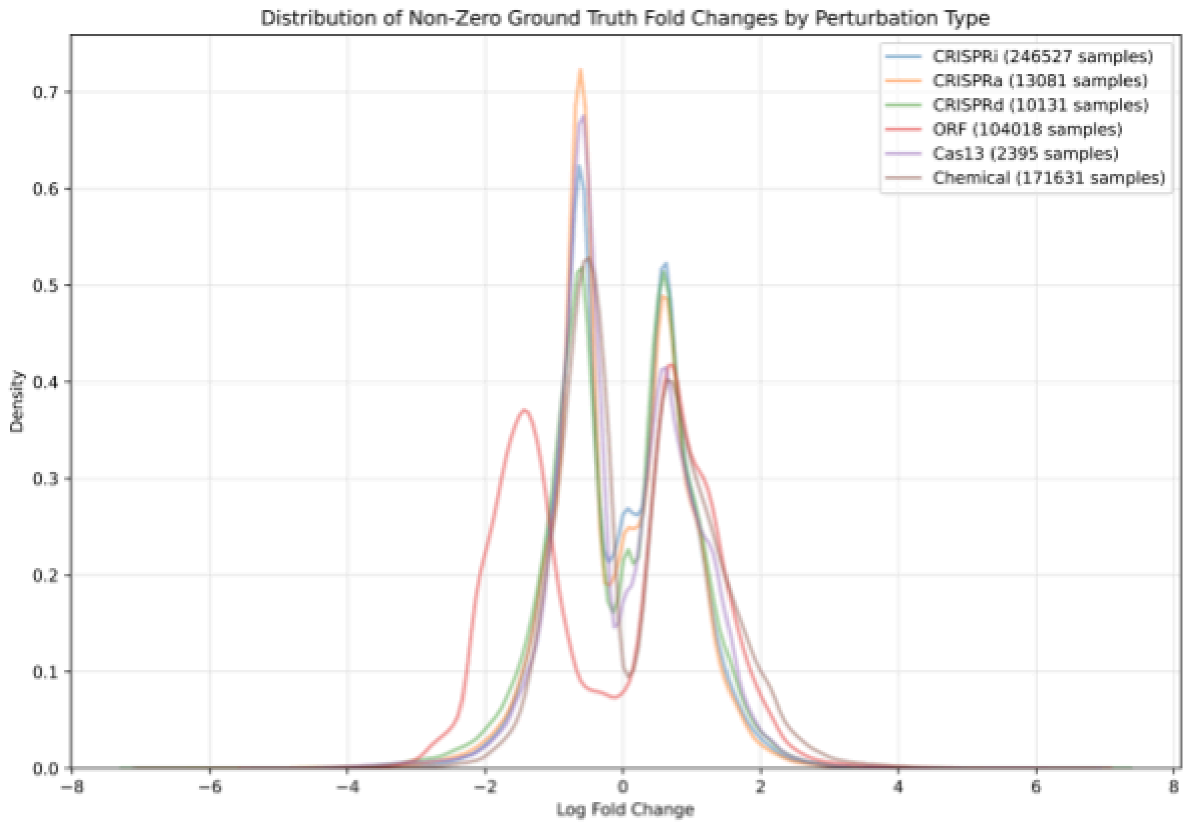
Distribution of Non-Zero Ground Truth Fold Changes by Perturbation Type
CRISPRi (246527 samples)
CRISPRa (13081 samples)
CRISPRd (10131 samples)
ORF (104018 samples)
Cas13 (2395 samples)
Chemical (171631 samples)
Density
Log Fold Change

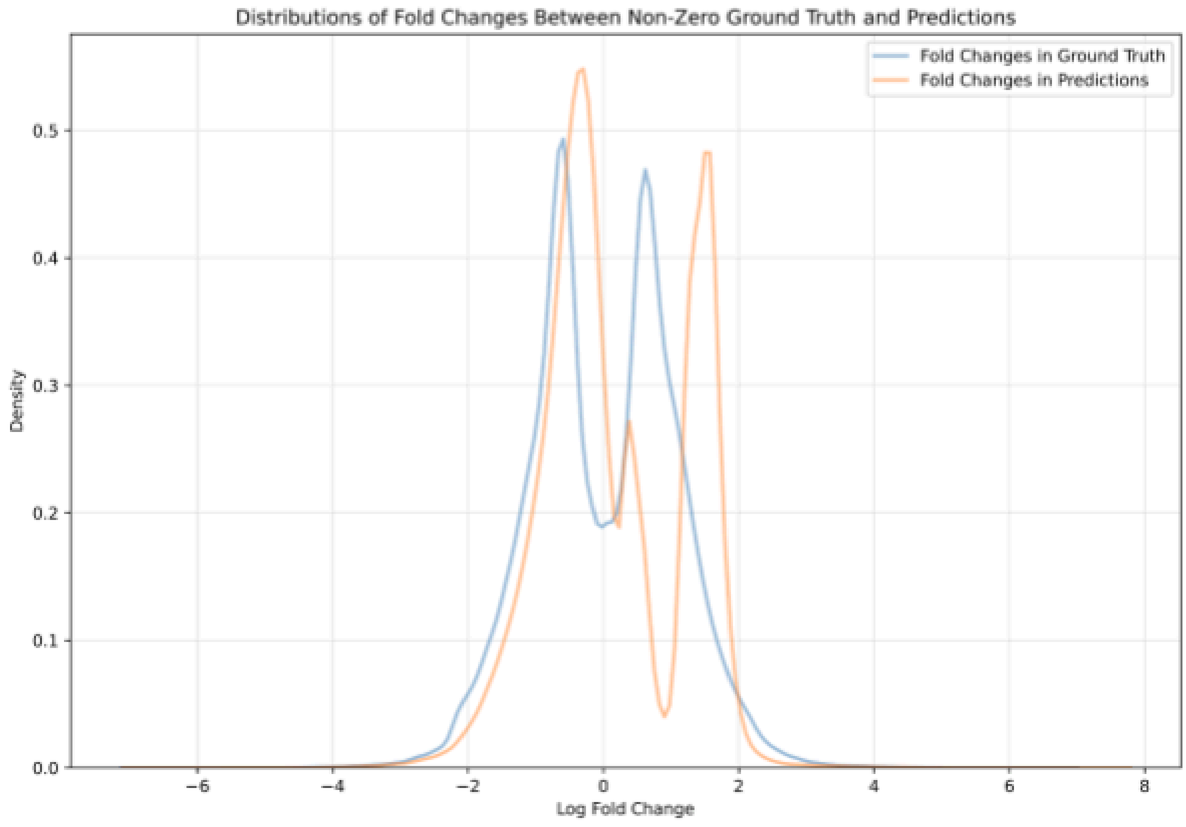
Distributions of Fold Changes Between Non-Zero Ground Truth and Predictions
Fold Changes in Ground Truth
Fold Changes in Predictions
Density
Log Fold Change

***Supplementary Figure 3.*** *Distribution of Fold Changes.*

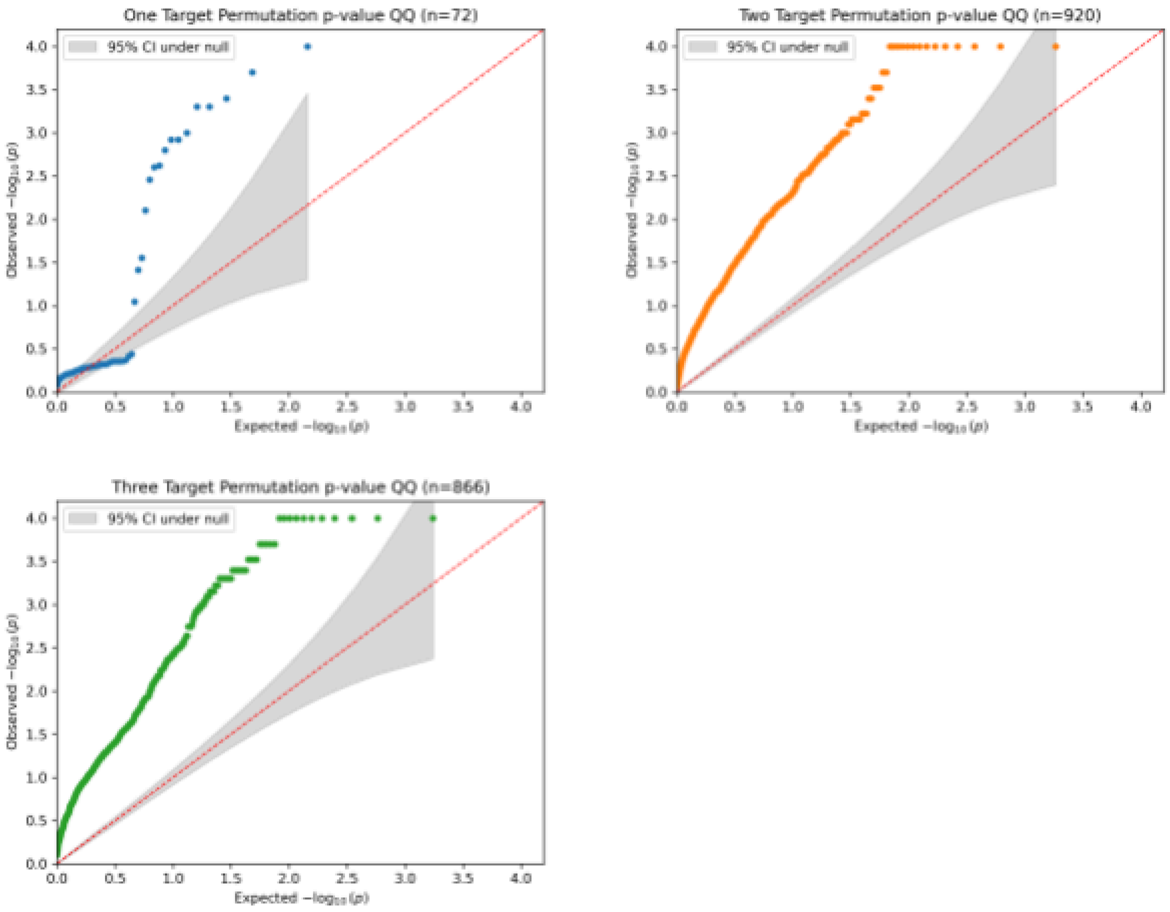


***Supplementary Figure 4.*** *Quantile-Quantile Plots negative logarithmic p-values of the permutation tests for genetic and chemical perturbation concordance of counterfactual predictions.*
